# Single cell clonal analysis identifies an AID-dependent pathway of plasma cell differentiation

**DOI:** 10.1101/2021.06.03.446763

**Authors:** Carmen Gómez-Escolar, Alvaro Serrano-Navarro, Alberto Benguria, Ana Dopazo, Fátima Sánchez-Cabo, Almudena R Ramiro

## Abstract

Germinal centers (GC) are microstructures where B cells that have been activated by antigen can improve the affinity of their B cell receptors and differentiate into memory B cells (MBCs) or antibody secreting plasma cells. Activation Induced Deaminase (AID) initiates antibody diversification in GCs by somatic hypermutation and class switch recombination. Here we have addressed the role of AID in the terminal differentiation of GC B cells by combining single cell transcriptome and immunoglobulin clonal analysis in a mouse model that traces AID-experienced cells. We identified 8 transcriptional clusters that include dark zone and light zone GC subsets, plasmablasts/plasma cells (PB), 4 subsets of MBCs and a novel prePB subset, which shares the strongest clonal relationships with PBs. Mice lacking AID have various alterations in the size and expression profiles of these transcriptional clusters. We find that AID deficiency leads to a reduced proportion of prePB cells and severely impairs transitions between the prePB and the PB subsets. Thus, AID shapes the differentiation fate of GC B cells by enabling PB generation from a prePB state.

## INTRODUCTION

During the immune response, B cells that have been stimulated by antigen with T cell help can engage in the germinal center (GC) reaction, where they can differentiate into either memory B cells (MBC) or high affinity, long-lived plasma cells (PC). GCs are key to the efficiency of the immune response and underlie the mechanism of action of most vaccination strategies. In GCs, B cells proliferate, modify their immunoglobulin genes by somatic hypermutation (SHM) and class switch recombination (CSR), are selected by affinity maturation, and terminally differentiate into alternate fates (Allen et al., 2007, Laidlaw and Cyster, 2020, Mesin et al., 2016, Shlomchik et al., 2019, Victora and Nussenzweig, 2012).

Activation induced deaminase (AID) initiates SHM and CSR (Muramatsu et al., 2000, Revy et al., 2000) with the deamination of cytosines on the DNA of immunoglobulin genes, which can be subsequently processed by various molecular pathways (Methot and Di Noia, 2017). In the case of SHM, AID deaminates the antigen recognizing, variable region of immunoglobulin genes, generating mutations which give rise to variants with altered affinity for antigen (Methot and Di Noia, 2017). In CSR, AID deaminations promote a recombination reaction between switch regions -highly repetitive sequences that precede constant regions-thus promoting the exchange of IgM/IgD isotypes by IgG, IgE or IgA isotypes encoded by downstream constant genes at the immunoglobulin heavy (IgH) locus (Methot and Di Noia, 2017). Therefore, immunoglobulin diversification by AID is central to the GC reaction.

GCs comprise two different compartments, the dark zone (DZ), where B cells proliferate and undergo SHM in their variable genes, and the light zone (LZ), where those B cells in which SHM has resulted in a BCR with increased affinity for the antigen, are selected in the context of T follicular helper cells and follicular dendritic cells (Allen et al., 2007, Victora and Nussenzweig, 2012). CSR takes place at the LZ, but it can also occur outside of the GC (Roco et al., 2019). B cells perfection affinity maturation with iterative cycles of mutation and proliferation in the DZ and positive selection in the LZ (Victora and Nussenzweig, 2012, Victora et al., 2010).

Notably, differentiation into the MBC and the PC fates from the GC is not stochastic; instead, higher affinity B cells preferentially differentiate into PCs, while MBCs generally show lower affinity for antigen (Mesin et al., 2016, Taylor et al., 2015, Kräutler et al., 2017, Phan et al., 2006, Shinnakasu et al., 2016, Suan et al., 2017, Viant et al., 2020). Accordingly, the frequency of SHM is on average higher in PCs than in MBCs (Laidlaw et al., 2020, Shinnakasu et al., 2016, Weisel et al., 2016). Likewise, BCR isotype influences the outcome of GC differentiation towards the PC or MBC fates (Gitlin et al., 2016, King et al., 2021, Kometani et al., 2013). This skewed selection into the MBC versus PC fate ensures high affinity protection by the PC effector compartment, while preserving an MBC reservoir of B cells with looser affinity which could be critical to provide a rapid defense against closely-related pathogens, as previously proposed (Kaji et al., 2012, Viant et al., 2020).

AID deficiency does not only promote a complete block in CSR and SHM, it also causes lymphoid hyperplasia, both in mouse and man (Muramatsu et al., 2000, Revy et al., 2000), indicating a role of AID in B cell homeostasis. Indeed, AID−/− mice have increased number of GC B cells upon immunization (Muramatsu et al., 2000), and AID−/− GC B cells show reduced apoptosis rate (Zaheen et al., 2009). Interestingly, AID−/− GC B cells tend to accumulate in the LZ and do not efficiently form PCs in mixed bone marrow chimeras (Boulianne et al., 2013). However, the contribution of AID to shaping B cell fate in GCs is not entirely understood. Moreover, the role of AID in GC clonal expansion and differentiation pathways has not been directly addressed.

Here, we have approached this question by combining single cell transcriptome analysis and V(D)J analysis of B cells from wild type and AID deficient mice. To that end, we have made use of a genetic mouse model that irreversibly labels cells that have expressed AID. We found that AID-experienced B cells clustered into 8 distinct transcriptional clusters, including LZ and DZ GC, 4 MBC subsets, a plasmablast PB/PC (PB) cluster and a novel prePB cluster, which shares strong clonal relationship with PB. The GC response in AID deficient mice showed alterations in cluster proportions as well as transcriptome differences in some of these clusters. Further, clonal relationships were profoundly altered in AID deficient mice, where the connection between prePB and PB clusters is severely impaired. Thus, our data reveal a critical role of AID in shaping the ultimate fate of B cell differentiation in GCs.

## RESULTS

### Single cell analysis of the GC response identifies 8 transcriptional clusters

To map GC differentiation at the single cell level, we made use of the Aicda-Cre^+/ki^; R26tdTom^+/ki^ (hereafter Aicda^Cre/+^) mouse model, which allows genetic tracing of cells that have expressed AID. In this model, the cDNA coding for the Tomato (Tom) fluorescent protein is inserted in the Rosa26 endogenous locus preceded by a transcriptional stop sequence that is flanked by loxP sites (R26tdTom allele). The Cre recombinase is inserted in the endogenous Aicda locus (*Aicda-Cre* allele) (Robbiani et al., 2008, Rommel et al., 2013). In Aicda^Cre/+^ mice, activation of the *Aicda* locus promotes Cre expression and excision of the transcriptional stop at the R26tdTom allele, unleashing the expression of the Tom protein. Thus, B cells that have been activated for AID expression, become irreversibly Tom+ (Figure S1A). To trigger a GC immune response, we first adoptively transferred CD4+ T cells from OT-II mice, which harbor a TCR recognizing a peptide from the ovalbumin (OVA) protein, into Aicda^Cre/+^ mice. Mice were immunized with OVA 1 day and 15 days after the transfer of OT-II cells. As immunization controls, we used OT-II transferred Aicda^Cre/+^ mice injected with PBS. Mice were sacrificed for analysis 15 days after the second immunization (Figure 1A). Flow cytometry analysis showed that OVA immunization expectedly resulted in the generation of Tom+ cells, which comprised GC B cells (GC, Tom+GL7+), plasmablasts and PCs (PB, Tom+CD138+) and putative memory B cells (pMem, Tom+ GL7-CD138−) (Figures 1B and 1C).

**Figure 1.**
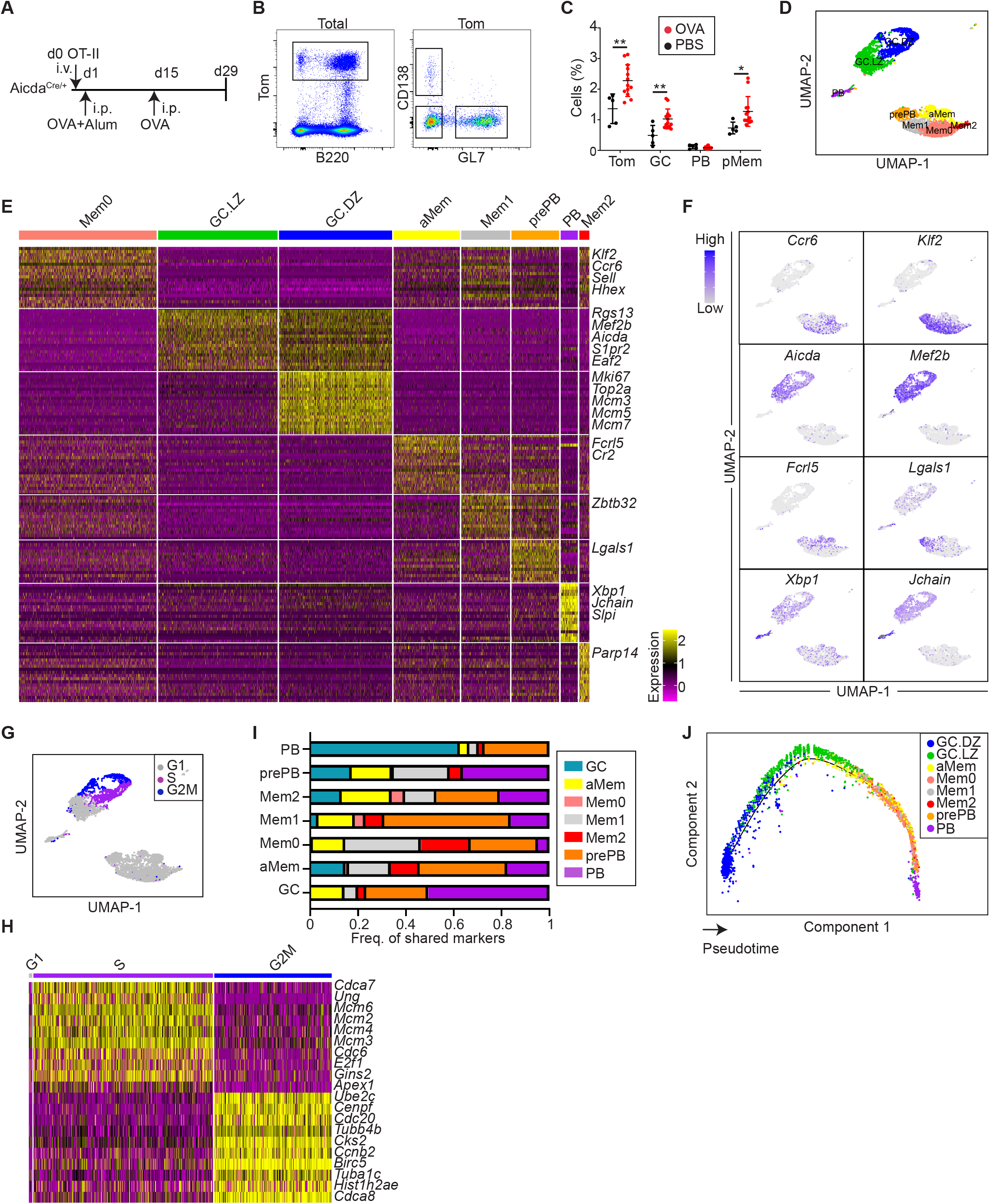
Single cell RNA sequencing of AID-labeled Tomato+ (Tom+) cells identifies eight cell clusters. **A.** Immunization protocol. Aicda^Cre/+^ mice were immunized intraperitoneally (i.p.) with OVA in alum (n=13) or with PBS (n=5) one day after OT-II CD4+ T cell transfer. Two weeks later, mice were boosted with OVA i.p. **B.** Representative flow cytometry plots of spleen Tom+ cells, germinal center B cells (GC; B220+ Tom+ GL7+), plasma cells/plasmablasts (PB; Tom+ CD138+) and putative memory B cells (pMem; Tom+ GL7− CD138−). **C.** Quantification of flow cytometry analysis of immunized mice as shown in A and B (statistics were calculated with unpaired t test comparing PBS (n=5) and OVA-immunized mice (n=13). **D.** Splenic Tom+ cells from two immunized Aicda^Cre/+^ mice were analyzed by single cell RNA sequencing (scRNA-seq) using the 10x Genomics platform. Cells were clustered based on transcriptomic data and mapped to an UMAP plot. Clusters are labeled as dark zone GC (GC.DZ), light zone GC (GC.LZ), plasmablast/plasma cells (PB), pre-plasmablast (prePB), Memory 0, 1 and 2 (Mem0, Mem1, Mem2) and atypical Memory (aMem). **E.** Heatmap showing expression of the top 15 upregulated genes within the clusters identified in C. Yellow indicates higher gene expression. Representative gene names are indicated on the right. **F.** UMAP plots showing expression of representative genes of the different B cells clusters, as shown in D. Blue color indicates higher gene expression. **G.** UMAP plot showing the cell cycle phase of individual cells in the different clusters as shown in D. **H.** Heatmap of G1, S and G2M subclusters of GC.DZ cells showing expression of the top 10 upregulated genes in S and G2M phases. **I.** Bar plot depicting proportions of markers with highest expression shared by each cluster with all the rest, as analyzed with QuickMarker. **J.** Monocle pseudotime analysis of the cells displayed in D. The projection is colored by cluster identity and the cells are ordered by Pseudotime. Error bars indicate mean ± standard deviation. Statistics were calculated with unpaired t test *P≤0.05, **P<0.01, ***P<0.001 and ****P<0.0001.

To analyze the B cell immune response at the single cell level we performed 10x Genomics analysis in Tom+ spleen cells isolated from OT-II transferred Aicda^Cre/+^ mice 15 days after the boost OVA immunization (Figure 1A). Two individual immunized mice were multiplexed by hashtag labeling (HTO, see materials and methods) and we performed gene expression and V(D)J sequencing of individual cells (Figure S1B). Seurat clustering of gene expression sequencing in individual Tom+ cells identified 8 independent transcriptional clusters (Figures S2A-C, Figure 1D, Table S1). GC B cells expectedly showed high levels of *Aicda*, *S1pr2* or *Mef2b* and distinctly comprised LZ (GC.LZ) and DZ (GC.DZ) clusters, as defined before (Victora et al., 2012, Victora et al., 2010) (Figures 1E and 1F, Figure S2D, Table S1). We found that the vast majority of proliferating Tom+ cells were contained in the GC.DZ cluster, and conversely, virtually all the cells (99%, 811/820 cells) in the GC.DZ cluster were in the S+G2M phases of the cell cycle (Figures 1G and 1H). Interestingly, UMAP projection precisely distinguished between cells in the S phase expressing high levels of replication genes (*Mcm2, Mcm3, Mcm4, Mcm6, Cdc6*, etc) and cells in the G2 and M phases, with high expression of mitotic genes (*Cdc20, Ccnb2, Cdca8*, etc) (Figures 1G and 1H). The PB cluster displayed high levels of *Xbp1*, *Jchain* and immunoglobulin genes (Figures 1E and 1F, Table S1) and was enriched in cells expressing the PB differentiation program (Figure S2E), as previously defined (Lin et al., 2003, Minnich et al., 2016).

Seurat gene expression analysis also detected a cluster of cells showing high expression of *Klf2* and *Ccr6* (Figure 1F, Table S1), previously associated with the MBC transcriptional program (Laidlaw et al., 2020, Suan et al., 2017), which we initially labeled as Mem cells (Figure S2A). Sub-clusterization of the Mem population identified three transcriptionally distinct MBC populations: one major subset (Mem0) with highest expression levels of *Klf2*, *Ccr6* and *Hhex*, one subset (Mem1) with highest levels of *Zbtb32* and *Vim*, and a minor subset (Mem2) with highest levels of *Irf7* and *Isg15* (Figures S2B and S2C, Figures 1D and 1E, Table S1). One additional distinct cluster (aMem) expressed high levels of *S1pr1*, *Ccr6* and *Cd38*, also consistent with a MBC identity (Kräutler et al., 2017, Laidlaw et al., 2020, Suan et al., 2017), as well as *Fcrl5*, recently associated with a subset of the so-called atypical MBC (Kim et al., 2019) (Figures 1D and 1E, Table S1). Finally, an additional cluster was identified (Figures 1D and 1E, Table S1), which displayed similarities with the PC and PB Immunological Genome Project gene sets (Figure S2F) and showed the second highest enrichment score with a PB signature (Minnich et al., 2016) (Figure S2E); thus, it was labeled as prePB cluster.

To get insights into the inter-relationships among the identified transcriptional clusters we first performed an analysis of transcriptional transitions using QuickMarkers (Table S2). For each cluster we quantified the proportion of markers shared with the highest frequency by the rest of the clusters. We found that PB makers were most frequently found in GC and prePB clusters, suggesting a higher rate of transcriptional transitions among these clusters. In addition, Monocle pseudotime analysis of transcriptional changes confirmed our initial cluster assignment, with DZ and LZ immediately preceding the differentiation of aMem, Mem0, 1 and 2, and with prePB immediately preceding the PB state (Figures 1I and 1J).

Thus, single cell transcriptome analysis of a T-dependent immune response identifies 8 distinct GC-related populations, including GC.DZ, GC.LZ, PB, 4 clusters of MBC and prePB.

### Single cell clonal analysis of the GC response

To further characterize the identified transcriptional clusters, we first analyzed SHM in V(D)J transcripts of individual cells. We found that the mutation load at the IgH variable region widely varied across cells from different populations, with the highest mutation frequency found in both GC.DZ and GC.LZ B cells (Figures 2A-C, Figure S3A). We found that cells in the prePB and PB clusters had similar mutation frequencies (Figures 2A-C, Figure S3A). Finally, MBC clusters harbored the lowest mutation frequencies, in agreement with previous reports (Viant et al., 2020, Weisel et al., 2016), with the exception of Mem1, where mutations were significantly higher (Figures 2A-C, Figure S3A). The presence of mutations in these clusters confirms the MBC assignment based on transcriptome clustering. In addition, the distinct mutational load observed in different Mem clusters suggests they can represent functionally distinct differentiation states. CSR analysis on V(D)J transcripts generally mirrored the SHM results, with the highest proportion of isotype switched cells within the GC.DZ and GC.LZ clusters (Figures 2D and 2E, Figures S3B and S3C).

**Figure 2.**
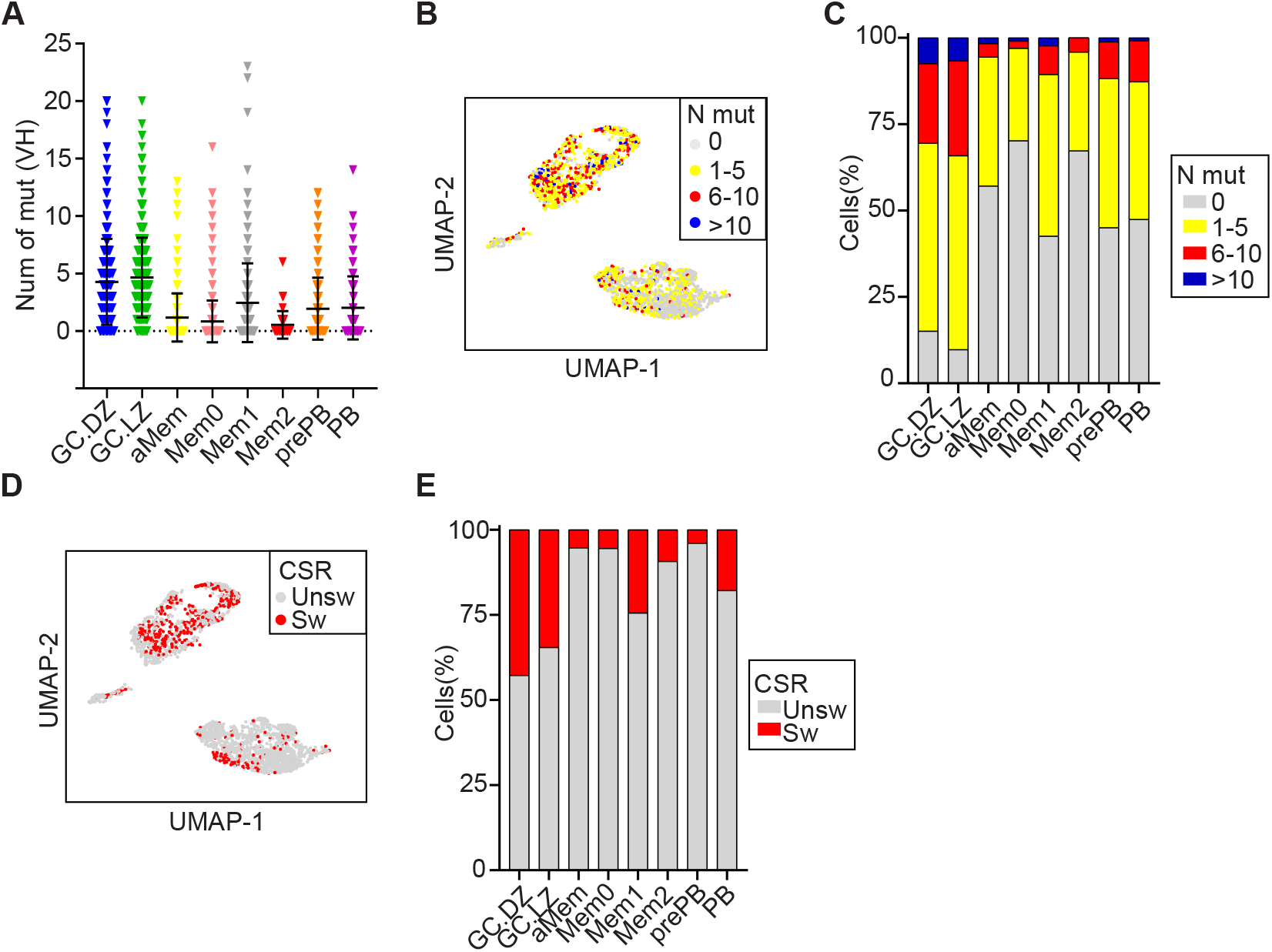
Single cell analysis of SHM and CSR during the GC reaction. **A.** Total number of somatic mutations at the IgH variable region (VH) in individual cells from the different B cell clusters. Symbols represent individual cells. Statistics were calculated with Kruskal-Wallis test using Dunn’s multiple comparison test (Figure S2A). **B.** UMAP plot showing mutational load in single cells of the clusters defined in Figure 1D. **C.** Quantification of data shown in B. **D.** UMAP plot showing CSR of individual cells (Unsw, unswitched; Sw, switched). **E.** Proportion of CSR in individual clusters (Unsw, unswitched; Sw, switched).

To identify clonal relationships among the different clusters we performed analysis of BCR sequences using the Immcantation pipeline. Cells were assigned to the same clone when they shared identical V(D)J segments and identical CDR3 lengths in both their IgH and IgL chains. We found that GC.DZ and GC.LZ clusters were highly enriched in clonally expanded cells (Figure 3A). Likewise, clonal sharing was most frequently observed between the GC.DZ and the GC.LZ clusters (Figures 3B and 3C, note that the circus plot represents all Mem clusters together for the sake of clarity, see Figure S4 for further details). These findings reflect the high proliferation rate at the DZ as well as the iterative transitions of B cells between the DZ and the LZ in GC. Our data also revealed clonal sharing among other clusters. For instance, clone 1 (containing 31 cells) was found several times in the GC.DZ and GC.LZ clusters, both in unswitched and in switched versions, but it was also found in the aMem and the prePB clusters (Figures 3D and 3E). In contrast, we found that clone 3 was shared between the aMem and the PB clusters (Figures 3D and 3E). Thus, clone 3 probably represents the end differentiation state of a clonal expansion that is already extinguished at the GC; in contrast, while clone 1 must have been at full expansion in the GC at the time of analysis, we can also identify cells that have transited to the aMem or the prePB clusters.

**Figure 3.**
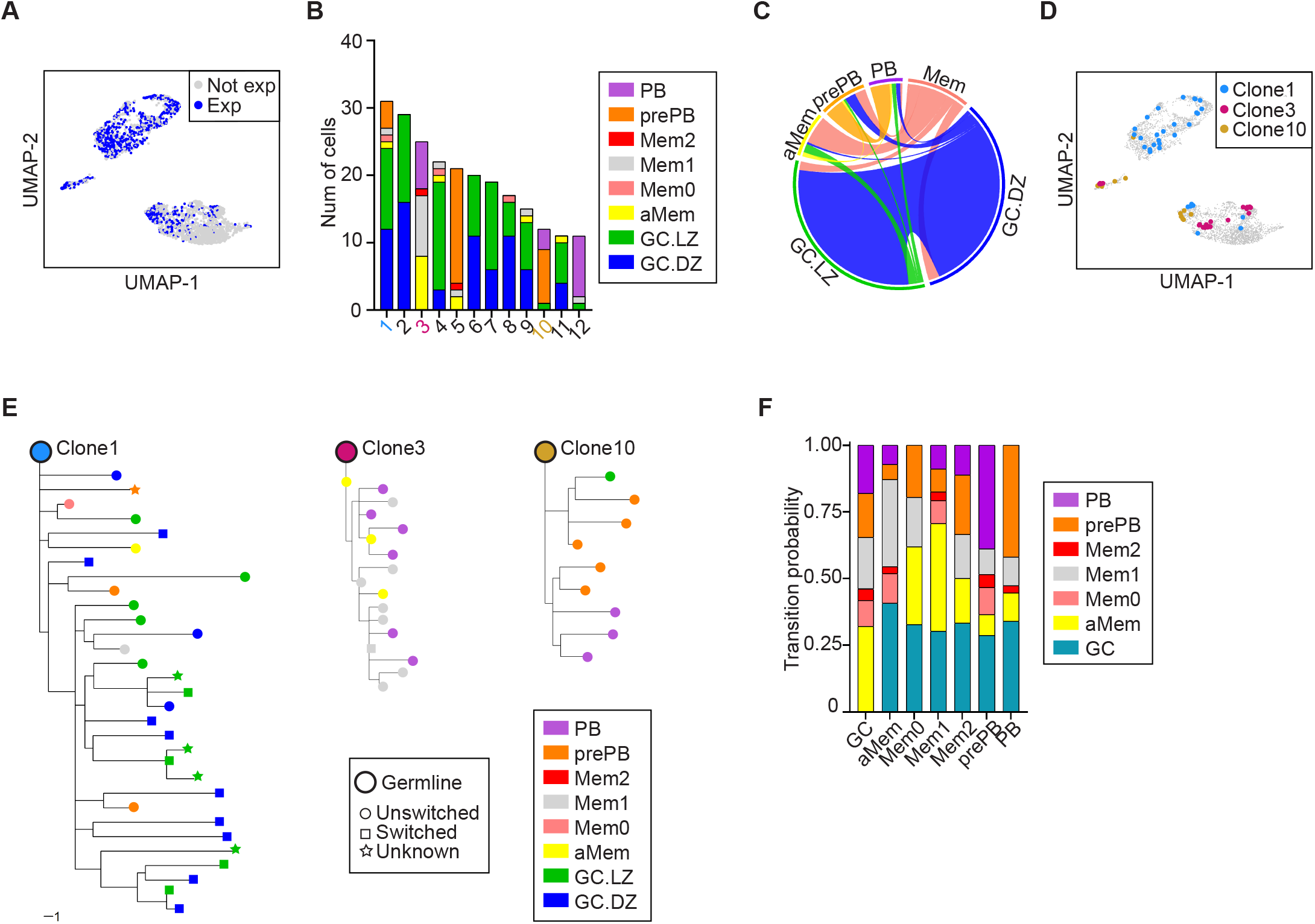
Single cell analysis of clonal relationships in GCs. **A.** UMAP plot showing the distribution among clusters of clonally expanded cells (Exp, expanded; Not exp, not expanded). **B.** Bar plot depicting the contribution of the different transcriptional clusters to expanded clones. Only clones >10 cells are shown. See Figure S3A and Table S3 for complete collection of clusters. **C.** Circos plot displaying pairwise clonal overlap between B cell clusters. Only clones shared between 2 clusters are shown. For the sake of clarity, Mem0, Mem1 and Mem2 clusters are shown together as Mem. **D.** UMAP plot showing 3 representative expanded clones (clone1, clone3, clone10). **E.** Trees showing phylogenetic relationships between IgH sequences of clone1, clone3 and clone10. **F.** Transition probabilities among the different clusters, based on the frequency of clonal sharing between 2 or more clusters.

V(D)J clonal sharing (Figure 3C) allows us to calculate the transition probabilities across differentiation states (Figure 3F, Table S3). We observed that the GC cluster interconnects with all other clusters, suggesting that GC cells can feed other subsets, and possibly also receive cells from the different subsets of MBC (Figures 3C-F). On the other hand, the frequent clonal sharing observed between Mem and aMem clusters suggests these clusters represent closely related states (Figure 3F). Interestingly, even though some clonal sharing is observed between Mem or GC and the PB cluster, the most predominant clonal relationship of PB is with the prePB cluster. This finding indicates that, while differentiation into the PB cluster can directly occur from the GC and Mem clusters, prePB is a very common intermediate state preceding the generation of PC/PB.

Thus, combined single cell transcriptome and V(D)J analysis of the GC response has allowed to establish clonal relationships between the GC clusters and all other clusters. Further, the newly identified prePB cluster shows the highest clonal sharing with PB, indicating that prePB cells represent a transition state preceding PB differentiation.

### Transcriptional clusters are shifted in AID deficient mice

To address the role of AID in GC differentiation at the single cell level, we generated Aicda-Cre^−/ki^; R26tdTom^+/ki^ mice (hereafter Aicda^Cre/−^ mice), where one *Aicda* allele is disrupted by the Cre recombinase and the other one is a knock-out allele (Figure S1A) (Muramatsu et al., 2000). Thus, AID deficient, GC-associated B cells can be genetically traced by the expression of the Tom protein. Control Aicda^Cre/+^ mice and AID deficient Aicda^Cre/−^ mice were immunized with OVA as above (Figure 1A) and the GC response in spleen was assessed by flow cytometry. We found that immunized Aicda^Cre/−^ spleens had a higher proportion of Tom+ cells, in agreement with the GC expansion found in AID deficient mice (Figure 4A) (Muramatsu et al., 2000, Zaheen et al., 2009). Within Tom+ cells, the proportions of GC and pMem were not altered, while the proportion of PB was severely reduced, consistently with previous reports (Figure 4B) (Boulianne et al., 2013). In addition, the DZ to LZ ratio was reduced in Aicda^Cre/−^ mice, as previously reported (Boulianne et al., 2013) (Figures 4C and 4D).

**Figure 4.**
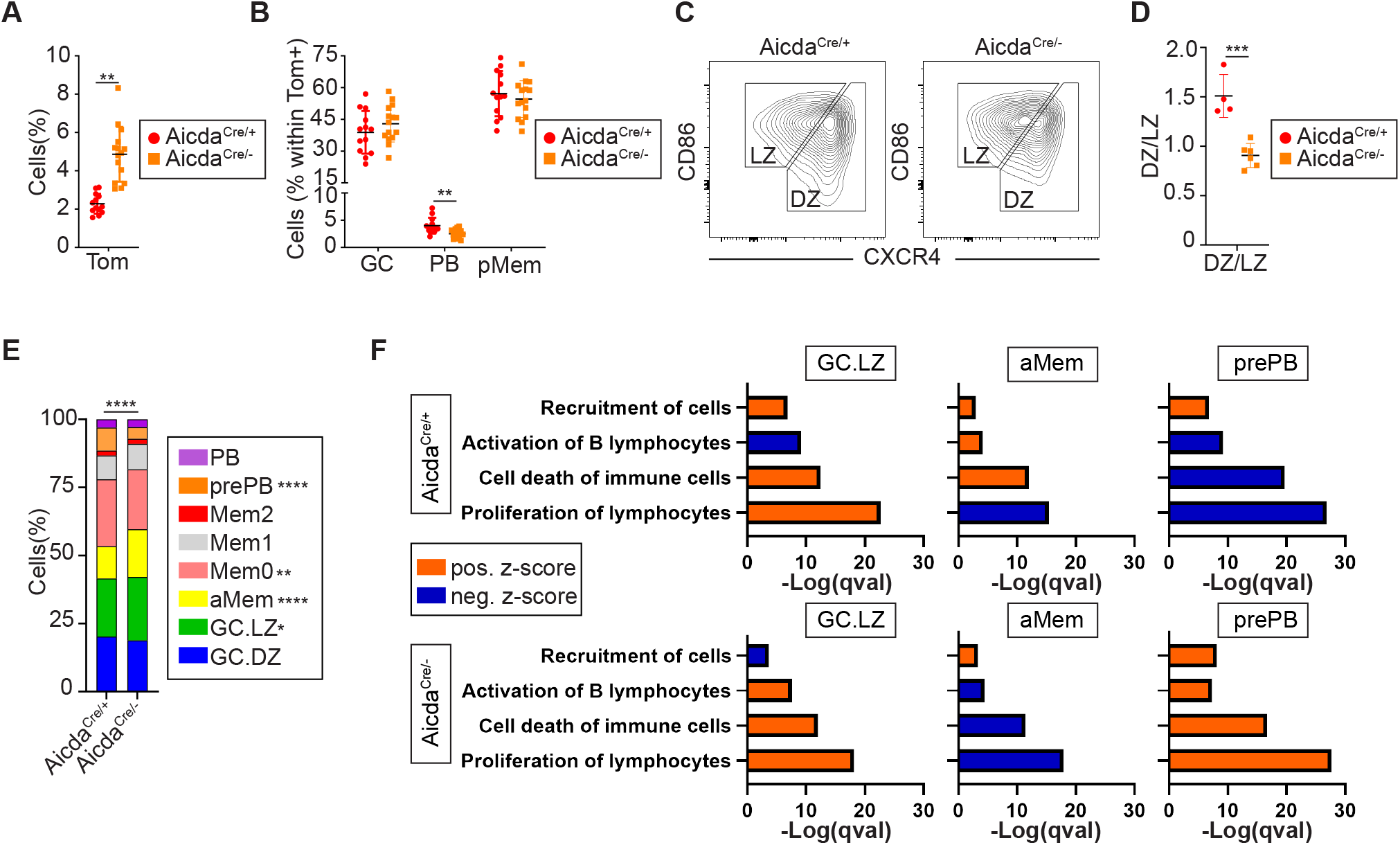
AID deficiency alters the proportions of GC-associated cell subsets. **A.** Aicda^Cre/−^ mice were immunized with OVA following the protocol in Figure 1A (n=15). Spleen Tom+ cells were quantified and compared to OVA-immunized Aicda^Cre/+^ mice. **B.** The proportion of GC B cells (B220+Tom+GL7+), PB (Tom+CD138+) and pMem (Tom+GL7-CD138−) was determined by flow cytometry within total Tom+ cells in Aicda^Cre/+^ (red symbols) and Aicda^Cre/−^ (orange symbols) mice. **C.** Representative flow cytometry plots and quantification **D.** of dark zone (DZ; B220+ Tom+ GL7+ CXCR4hi CD86lo) and light zone (LZ; B220+ Tom+ GL7+ CXCR4lo CD86hi) GC B cells. DZ/LZ ratio is depicted in the right panel for Aicda^Cre/+^ (n=4) and Aicda^Cre/−^ (n=6) mice immunized with OVA. **E.** Spleen Tom+ cells from two immunized Aicda^Cre/−^ mice were analyzed by scRNA-seq. Cluster labels from Aicda^Cre/+^ mice (Figure 1C) were transferred to Aicda^Cre/−^ cells. Bar plot showing the proportions of the different B cell clusters in Aicda^Cre/+^ and Aicda^Cre/−^. **F.** Comparative pathway enrichment analysis of differentially expressed genes (DEGs) in GC.LZ, aMem and prePB clusters in Aicda^Cre/−^ mice (orange symbol), compared to Aicda^Cre/+^ (red symbol) mice. Analysis was performed with Ingenuity Pathway Analysis (IPA). Terms are colored by z-score (activation status). Positive values of z-score indicate activation of the pathway, while negative values indicate pathway inhibition. Benjamini-Hochberg adjusted p-values (qval) are represented. Error bars indicate mean ± standard deviation (A, B, and D). Statistics were calculated with unpaired t test (A, B, and D) and Chi-square (E, all clusters) and Fisher test (E, individual clusters). *P≤0.05, **P<0.01, ***P<0.001 and ****P<0.0001.

To further characterize the GC disturbances in AID deficient mice, we performed single cell analysis of Tom+ cells using 10x Genomics. Transfer of the transcriptome data into the clusters defined for wild type B cells (as shown in Figure 1D) showed that Aicda^Cre/−^ mice had altered proportions of several subsets (Figure 4E). Specifically, in addition to confirming the increase in GC.LZ cells, we found that the proportion prePB was severely diminished and the proportion of aMem was increased in Aicda^Cre/−^ mice (Figure 4E). Changes in subset proportions were accompanied by transcriptional alterations in several biological pathways (Figure 4F). Notably, the apoptotic signaling pathway was reduced in aMem, but was increased in prePB cells (Figure 4F), suggesting that AID deficiency interferes with prePB survival and thus compromises the transition to PB differentiation. In line with this idea, pseudotime analysis of AID deficient cells revealed a branch point in a late GC state (Figure 5A) that corresponds with distinct cell fates (Figure 5B). Thus, the mainstream branch (A) leads to the generation of PB cells, while branch B fails to give rise to the PB state and is instead biased towards the MBC fate (Figure 5C). Together, these results indicate that AID deficiency has profound consequences in B cell fate and differentiation during the GC response, by compromising the transition to terminally differentiated PB cells from prePB cells and instead promoting the accumulation of MBCs. This is consistent with the finding that MBCs have low mutational load (Shinnakasu et al., 2016) (Figure 2A) and that low affinity GC B cells preferentially differentiate into MBCs (Viant et al., 2020).

**Figure 5.**
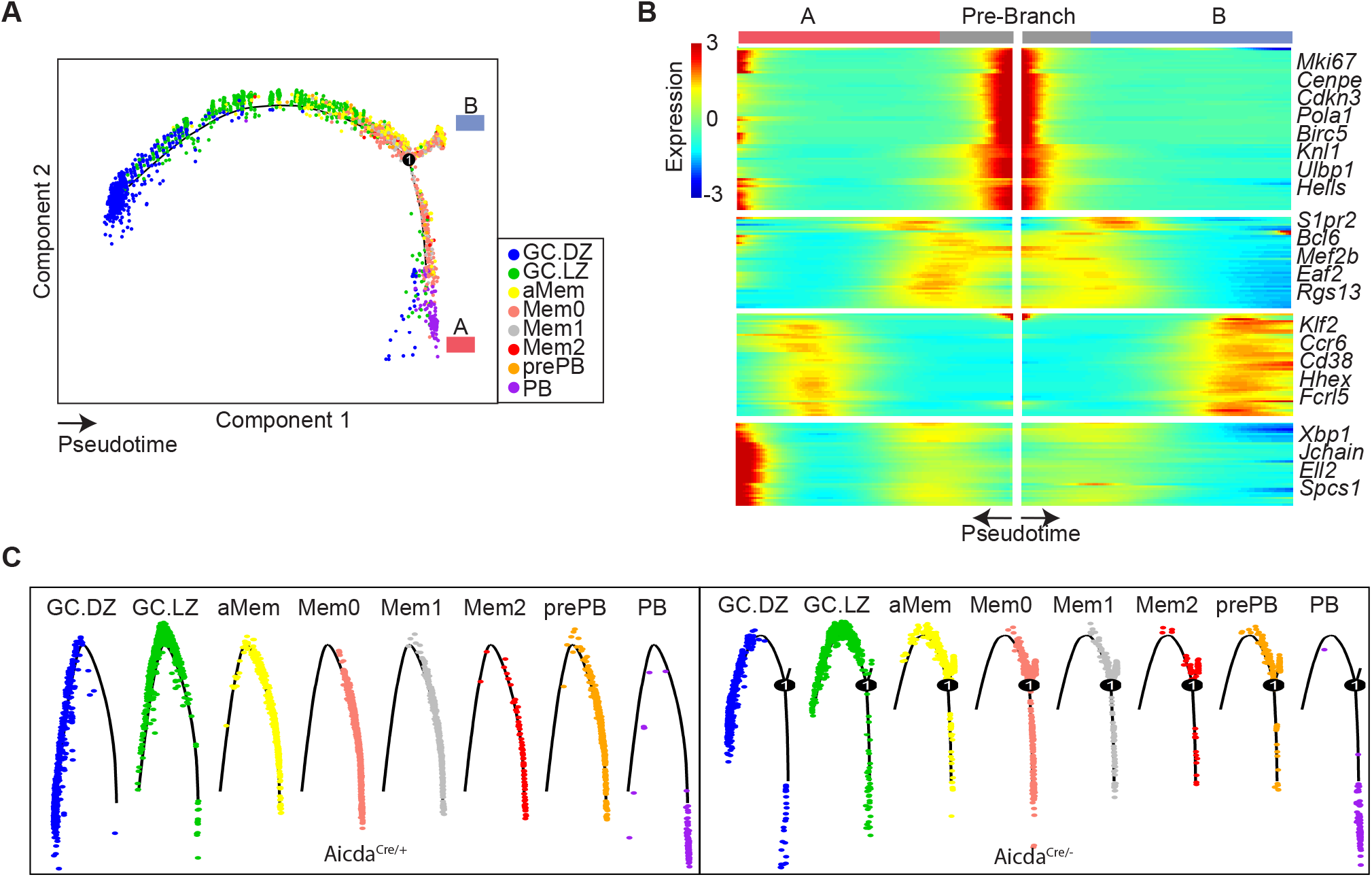
Pseudotime differentiation in AID deficient GCs. **A.** Monocle pseudotime analysis by of Aicda^Cre/−^ cells. The projection is colored by cluster identity. Number one in the circle identifies the branching point for branches A and B. **B.** Heatmap of pseudotime gene expression changes, showing representative DEGs between branches A and B. Genes are clustered hierarchically in modules with similar branch-dependent expression patterns. **C.** Mapping of cluster identities in the pseudotime plots of Aicda^Cre/+^ (left) and Aicda^Cre/−^ (right) cells.

### AID deficiency alters B cell differentiation during the GC response

To approach the consequences of AID deficiency in the clonal relationships and in the differentiation fate in GCs, we combined transcriptomics with V(D)J analysis in single cells. Clonal analysis of Tom+ cells showed that Aicda^Cre/−^ mice had more and larger expanded clones (Figures 6A and 6B). Increased clonal expansion in Aicda^Cre/−^ mice was not common to all cell clusters; rather, it was specifically observed in GC.DZ, aMem, Mem0 and Mem1 clusters (Figure 6C), indicating that AID can impact on proliferation, survival or differentiation at distinct stages of the GC reaction, in agreement with our transcriptome data. Analysis of clonal sharing between different clusters in Aicda^Cre/−^ mice expectedly showed the greatest interconnection between GC.DZ and GC.LZ clusters (Figure 6D, Figure S4A), similarly to our findings in Aicda^Cre/+^ mice (Figure 3B, Figure S4A). However, a closer look at pairwise clonal relationships between clusters showed major alterations in Aicda^Cre/−^ mice compared to Aicda^Cre/+^ mice (Figure 7A). Most notably, the clonal relationships of prePB and PB cells were severely reduced in Aicda^Cre/−^ mice (Figure 7A). To specifically quantify these alterations, we used clonal sharing frequencies (Table S3) to calculate the transition probabilities between any given cluster and all the rest and compared Aicda^Cre/−^ with Aicda^Cre/+^ mice (Figures 7B and 7C, Figure S4C). We found that, in Aicda^Cre/−^ mice, the prePB to PB transition is very much reduced and instead, the transition probability between aMem and PB is enhanced. These results indicate that, in the absence of AID, differentiation into PB is partially compromised due to a block at the prePB state, and this deficiency is partially compensated by an increased differentiation of PB from the aMem state.

**Figure 6.**
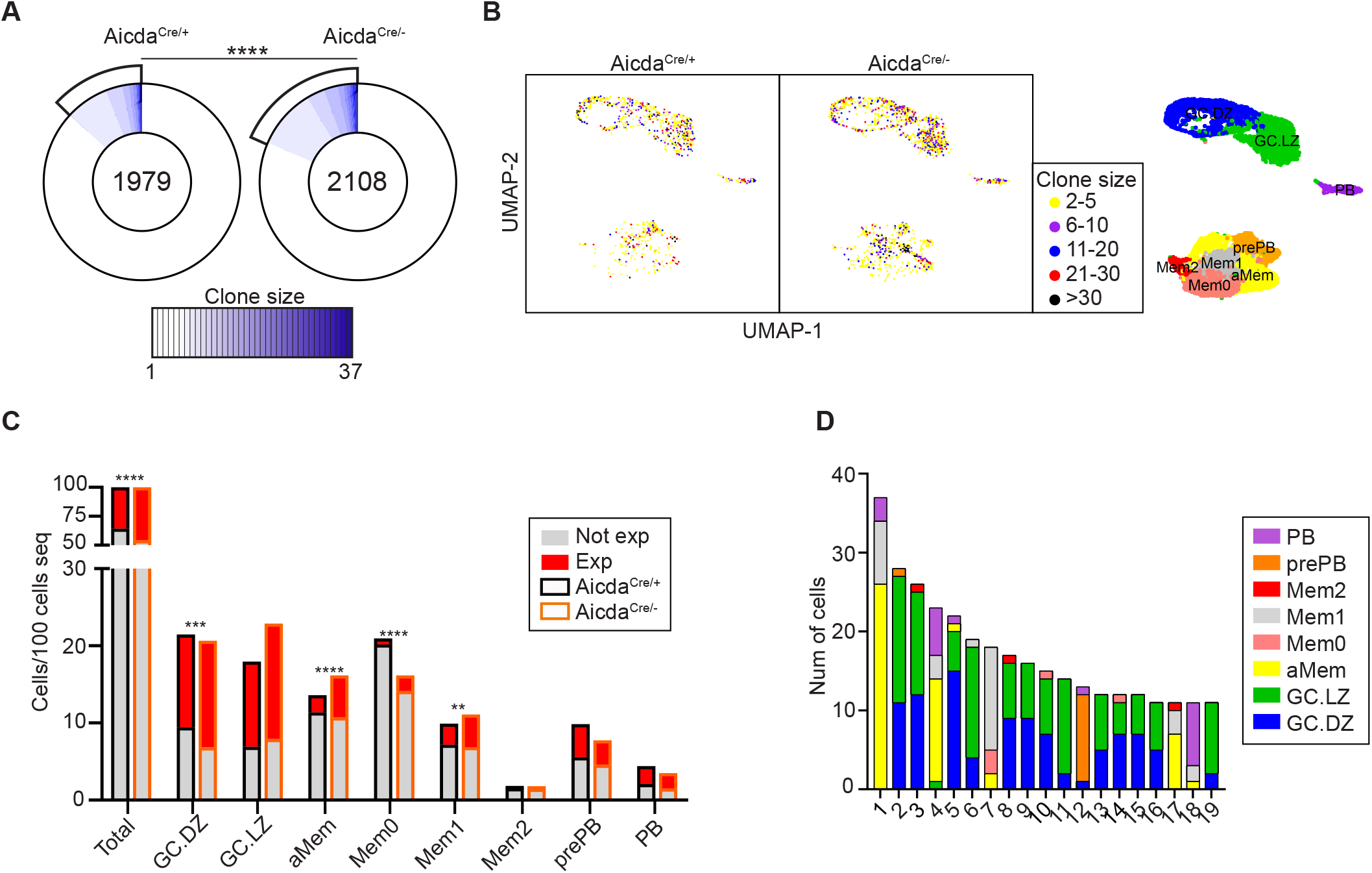
Increased expansion of B cell clones in AID deficient mice. **A.** Pie charts depicting the clonal size of Tom+ cells from Aicda^Cre/+^ and Aicda^Cre/−^ immunized mice. Clones are defined as cells sharing identical IGH and IGL V and J sequences and CDR3 lengths. Number in the inner circle indicates the total number of unique sequences. White represents sequences found only once and blue gradient represents increasing clonal sizes. Pie sector sizes are proportional to clone sizes. Two-sided Fisher’s exact test was used to compare the proportions of expanded and non-expanded clones. **B.** UMAP plot showing clone sizes in Aicda^Cre/+^ and Aicda^Cre/−^ mice. Cells are colored according to the size of the clone they belong to. Unique events are not shown. For reference, UMAP plot of transcriptional clusters is shown on the right. **C.** Clonal expansion in transcriptional clusters. Quantification of expanded and non-expanded cells is shown for individual clusters, normalized to the total number of cells sequenced per genotype. Two-sided Fisher’s exact test was used to compare the proportions of expanded versus non-expanded clones in each cluster. **D.** Bar plot depicting the contribution of the different transcriptional clusters to Aicda^Cre/−^ expanded clones. For simplicity, only clones >10 cells are shown. See Figure S3A and Table S3 for complete lists of clones.

**Figure 7.**
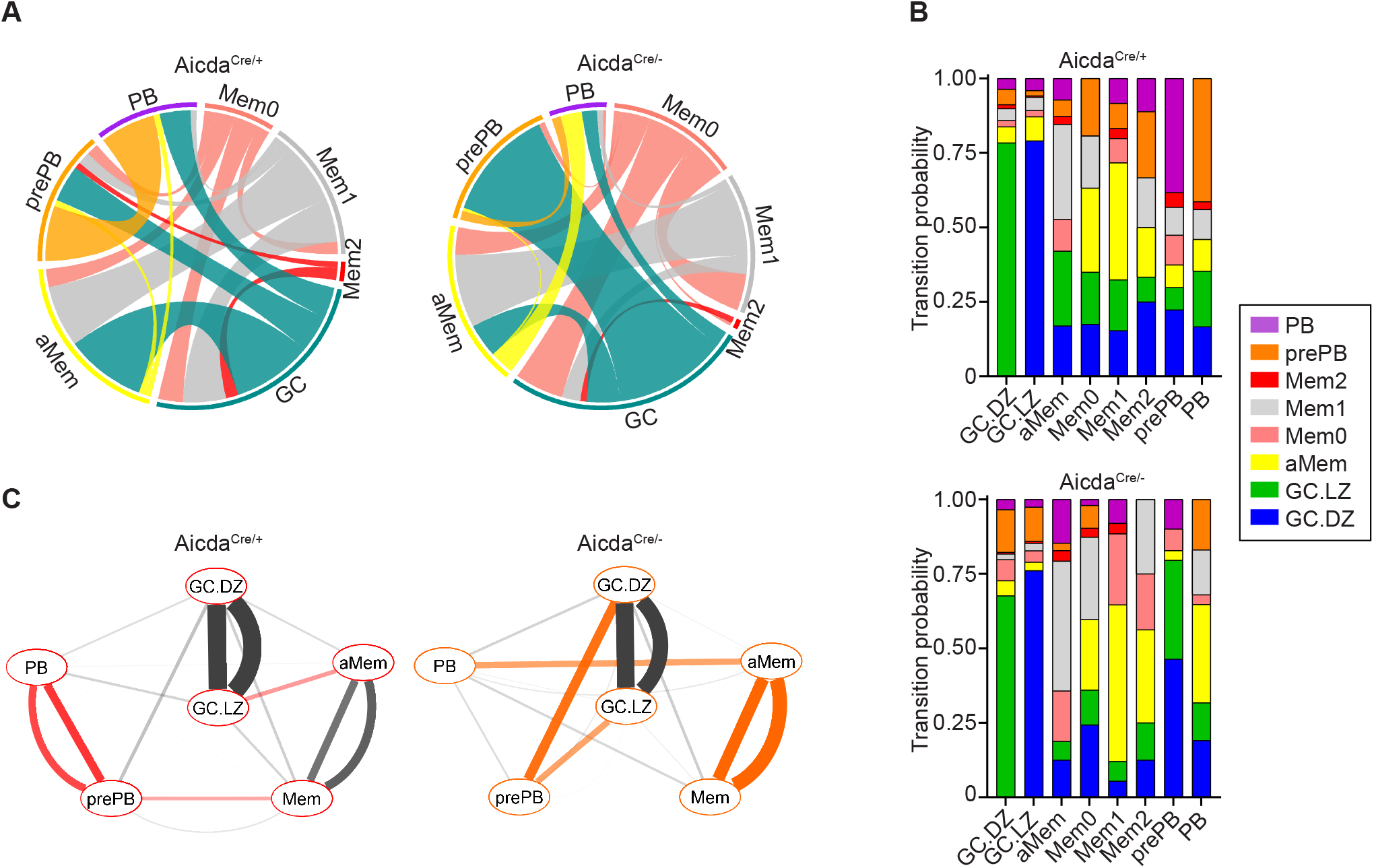
Altered B cell fate decisions in AID deficient mice. **A.** Circos plots showing pairwise clonal sharing between B cell clusters in Aicda^Cre/+^ and Aicda^Cre/−^ mice. GC.LZ and GC.DZ populations were grouped and shown as GC for the sake of simplicity. Complete clonal sharing relationships are shown in Fig S4B. **B.** Transition probabilities among the different clusters in Aicda^Cre/+^ and Aicda^Cre/−^, calculated as the frequency of clonal sharing between 2 or more clusters. Statistics were calculated with Fisher test. *P≤0.05, **P<0.01, ***P<0.001 and ****P<0.0001. **C.** Cytoscape representation of cluster interactions in Aicda^Cre/+^ and Aicda^Cre/−^ immune response, based on their clonal sharing probabilities. For each pairwise transition two lines are shown, which depict the probabilities with respect to each cluster of the pair. Red and orange connecting lines show the most increased cluster relationships in Aicda^Cre/+^ and Aicda^Cre/−^ mice, respectively. Mem0, Mem1 and Mem2 clusters are shown together for simplicity. See Figure S4C for complete interactions.

## DISCUSSION

In this study we have examined the role of AID in GC differentiation by combining single cell transcriptome and V(D)J sequencing. Several recent papers have performed single cell analysis of GCs both in mouse (Laidlaw et al., 2020, Nakagawa et al., 2021, Riedel et al., 2020, Wong et al., 2020) and human (Holmes et al., 2020, King et al., 2021), employing distinct isolation strategies. Here we have made use for the first time of an AID-based genetic tracer as a proxy to isolate GC-derived cells. While this tracing approach can potentially include a small fraction of extrafollicular activated B cells, it provided the best strategy to precisely address the loss of AID in those cells that have been programmed for AID expression.

Our study has identified 8 distinct transcriptional clusters within AID labeled cells, with very accurate identification of the LZ and DZ compartments as previously described (Victora and Nussenzweig, 2012, Victora et al., 2010), as well as the plasmablast/PC subset. Interestingly, we found that GC B cells harbored the highest mutational load of all clusters, indicating that the most highly mutated GC B cells do not proceed to either the PC or the MBC fates and instead remain at the GC. This possibly reflects a fraction of the GC B cells in which excessive SHM is detrimental for antigen binding/affinity and thus will not be positively selected. In addition, we identified 4 Mem clusters, which indicates a heterogeneity within the MBC compartment that has been previously observed by others (Shlomchik et al., 2019), also in single cell sequencing analyses (King et al., 2021, Laidlaw et al., 2020, Riedel et al., 2020). Transcriptional assignment of MBC is particularly challenging due to the ample transcriptome overlap with naïve B cells; however, our assignment has been aided by the additional restriction of AID tracing as well as the detection of SHM at V(D)J. In this regard, the different mutational load found in the various Mem clusters could reflect different times of exposure to SHM at the GC and possibly functional specialization (Shlomchik et al., 2019). In addition, clonal sharing among different Mem clusters suggests there is certain plasticity across MBC subsets, an observation that credits further investigation. Interestingly, we identified a prePB transcriptional cluster that shared the largest clonal similarities with the PB cluster, thus indicating that prePB cells can be PB immediate precursors. Given that this prePB cluster does not show particularly high levels of *Prdm1*, *Xbp1* or immunoglobulin transcripts, we speculate that they could be early GC or MBC emigrants skewed to PB differentiation.

Here we have confirmed the finding that AID deficiency impairs PB/PC differentiation and decreases the DZ/LZ ratio at GC (Boulianne et al., 2013, Zaheen et al., 2009). In addition, our single cell analysis has shown that AID deficiency severely reduces the proportion of prePB cells while favoring the accumulation of aMem. These alterations underlie a complex differentiation shift in the GCs of AID deficient mice, which is revealed as changes in transcriptional programs as well as in clonal relationships. Of note, since AID itself is not a transcriptional regulator, it is safe to argue that the observed transcriptional changes in Aicda^Cre/−^ mice are a consequence of secondary immunoglobulin diversification mediated by AID. Interestingly, we observed that AID modulates B cell survival and proliferation at distinct stages of GC differentiation, probably reflecting changes in antibody affinity maturation and immunoglobulin isotype-dependent checkpoints.

Importantly, our study has established for the first time a direct link between these transcriptional shifts and the clonal expansion and relationships among different clusters. Thus, in the absence of AID, clonal sharing between prePB and PB clusters is very much reduced, while the more expanded aMem cluster shows increased clonal relationships with the PB cluster. This finding suggests that the differentiation into PB in AID deficient mice is partially compensated by an alternative differentiation track which, intriguingly, involves an MBC intermediate that in control mice only rarely seems to be a precursor of PB. While further research is needed to pinpoint the exact molecular mechanisms underlying our observations, this study provides the first single cell map on the role of AID in GC differentiation.

## ACKNOWLEDGMENTS

We thank all the members of the B lymphocyte Biology lab for helpful suggestions, Sonia Mur for technical assistance, Virginia G de Yebenes for critical reading of our manuscript, Julia Merkenschlager, Carlos Torroja and Enrique Vazquez for their advice on single cell sequencing and analysis, the CNIC Flow Cytometry for assistance on cell analysis and separation and the CNIC Genomics Unit for single cell sequencing. We also thank Sergio Roa, Alicia G Arroyo and David Sancho for kindly sharing mouse lines with us. CG-E is supported by a fellowship awarded by La Caixa España 2017 and AS-N is an FPI Severo Ochoa fellow (PRE2018-083475). AB, FS-C and ARR are supported by CNIC. This project was funded by grants from the Spanish Ministerio de Economía, Industria y Competitividad (SAF2016-75511-R), the Spanish Ministerio de Ciencia e Innovación (PID2019-106773RB-I00/ AEI / 10.13039/501100011033) and the “la Caixa” Banking Foundation under the project code HR17-00247 to ARR. FS-C received support from the Spanish Ministerio de Economía y Competitividad (RTI2018-102084-B-I00). The CNIC is supported by the Instituto de Salud Carlos III (ISCIII), the Ministerio de Ciencia e Innovación (MCIN) and the Pro CNIC Foundation, and is a Severo Ochoa Center of Excellence (SEV-2015-0505)

## AUTHOR CONTRIBUTIONS

A. Ramiro designed and supervised the study; C. Gómez-Escolar performed the experiments; A. Benguria and A. Dopazo performed RNA sequencing; A. Serrano-Navarro and C. Gómez-Escolar performed computational analyses; F. Sanchez-Cabo contributed to computational analyses. A. Ramiro and C. Gómez-Escolar analyzed data and prepared figures; A. Ramiro and C. Gómez-Escolar wrote the manuscript.

## METHODS

### Mice, cell transfer and immunization

Aicda-Cre^+/ki^ mice (Rommel et al., 2013) and Rosa26tdTomato^+/ki^ mice are from Jackson laboratories (Aicda^tm1(cre)Mnz^, 007770 and 007909). Aicda^Cre/+^ mice were generated by breeding Aicda-Cre^+/ki^ to Rosa26tdTomato^+/ki^ mice. C57BL/6-Tg (TcraTcrb) 425Cbn/J mice (OT-II) express a TCR specific for the OVA peptide (amino acid residues 323–339) (Jackson Laboratory 004194). Aicda^−/−^ mice were a generous gift of Dr Tasuku Honjo (Muramatsu et al., 2000). Aicda^Cre/−^ mice were generated by crossing Aicda-Cre^ki/ki^;Rosa26tdTomato^ki/ki^ mice with Aicda^+/−^ mice.

CD4^+^ OT-II spleen T cells were enriched by negative selection with the EasySep™ Mouse CD4^+^ T Cell Isolation Kit (STEMCELL) and 5·10^4^ cells were transferred intravenously to groups of 10 weeks-old Aicda^Cre/+^ and Aicda^Cre/−^ mice. 16 hours after the transference, mice were immunized intraperitoneally with 50 μg of OVA (Sigma-Aldrich) in PBS, precipitated in alum (Imject Alum, Thermo Scientific) in a ratio of 1:2:1. Booster immunization was performed two weeks later, by intraperitoneal injection of 50μg of OVA in PBS. Control mice were injected with PBS. Mice were euthanized two weeks after the second immunization.

Animals were housed in the Centro Nacional de Investigaciones Cardiovasculares animal facility under specific pathogen-free conditions. All animal procedures conformed to EU Directive 2010/63EU and Recommendation 2007/526/EC regarding the protection of animals used for experimental and other scientific purposes, enforced in Spanish law under RD 53/2013. The procedures have been reviewed by the Institutional Animal Care and Use Committee (IACUC) of Centro Nacional de Investigaciones Cardiovasculares, and approved by Consejeria de Medio Ambiente, Administración Local y Ordenación del Territorio of Comunidad de Madrid

### Flow cytometry

Single cell suspensions were obtained from spleens and erythrocytes were lysed (ACK Lysing Buffer). Fc receptors were blocked with anti-mouse CD16/CD32 antibodies (BD Pharmingen™) and cells were stained with fluorophore or biotin-conjugated anti-mouse antibodies to detect B220 (RA3-6B2), GL7 (GL7), CD138 (281-2), CD38 (HIT2), CXCR4 (2B11) and CD86 (GL1). Streptavidin (BD) was used for biotin-conjugated antibodies. Live cells were detected by 7AAD (BD Pharmingen) or LIVE/DEAD Fixable Yellow Dead Cell Stain (Thermo Fisher) staining. Samples were acquired on LSRFortessa and analyzed with FlowJo V10.4.2 software.

### Cell sorting and scRNA-seq

Four different TotalSeq™-C antibodies were used for Hash Tag Oligonucleotide (HTO) labelling before cell sorting (Biolegend, M1/42M30-F11) following the manufacturer’s standard protocol. HTO stained Tom+ spleen cells were purified from immunized mice using Sy3200 Cell Sorter (SonyBiotechnology). After sorting, Tom+ cells from the four mice were pooled together and diluted at a concentration of 1,500 cells/μl. 17,500 cells were used for the Chromium 10X Genomics System following the manufacturer’s protocol (Chromium Single Cell 5’ Reagent Kit, v1.1 Chemistry). Single cell cDNA was separated into three aliquots for HTO, scRNA-seq and V(D)J-enriched library generation from the same input sample. The three libraries were sequenced with the HiSeq4000 System (Illumina).

### scRNA-seq data analysis

Cell Ranger mouse mm10 genome reference (v3.0.0 November 18, 2018) was used for alignment of FASTQ files using 10x Genomics Cell Ranger software v3.1.0. A unique count table of gene expression for the four samples was generated. HTO antibodies were used for sample demultiplexing using Seurat R package(Butler et al., 2018) (v3.1.5). An average of 57,343 reads per cell and a median value of 1,748 expressed genes per cell were obtained. The estimated number of total cells recovered was 10,454. Mice were grouped by genotype for further analyses. Single cell gene expression analysis was performed with Seurat. UMI counts measuring gene expression were log-normalized and cells with mitochondrial reads greater than 8% of all transcripts were removed. A small number of non-B cells were identified and filtered out. After quality filtering, there were a total of 8,116 B cells from the 4 samples. The top 2,000 variable genes were identified with the “FindVariableFeatures” function and scaled. UMAP projection plots were generated using the top 30 principal components. Clusters were identified with “FindClusters” function with a resolution of 0.25. Upregulated marker genes for each cluster were inferred with “FindAllMarkers” function (Wilcoxon test), using normalized expression values. Top 15 markers per cluster were visualized using “DoHeatmap” function. Cluster identities were assigned based on the expression of markers for different B cell types and using “MyGeneSet” tool from the Immunological Genome Project Databrowser. Markers used for assignment of cluster identities included: *Aicda*, *Fas*, *Mef2b*, *Bcl6*, *S1pr2* and *Rgs13* for GC B cells; *Klf2*, *Ccr6*, *Sell*, *Zbtb32* and *Hhex* for MBCs, *Xbp1*, *Scd1*, *Jchain* and *Slpi* for PBs; *Fcrl5*+ memory B cell signature(Kim et al., 2019) for aMem. Signature enrichment was assessed using the “AddModuleScore” function in Seurat package. Further subclustering of the “Mem” cluster was performed and applied to main UMAP projection. Cell cycle scores were calculated using the “CellCycleScoring” function in the Seurat package using previously defined gene signatures for each cell cycle stage. Second best hit marker analysis was performed using the function “QuickMarkers” in the SoupX package(Young and Behjati, 2020), with a gene frequency threshold of 10%.

Ingenuity Pathway Analysis (IPA) was used the identify the distinct biological functions altered between Aicda^Cre/+^ and Aicda^Cre/−^ within the specified transcriptomic clusters.

### Pseudotime analysis

Pseudotime analysis was performed using the R package Monocle 2.16.0 (Trapnell et al., 2014). A single cell trajectory was constructed by Discriminative Dimensionality Reduction with Trees (DDRTree) algorithm using all genes differentially expressed among Seurat clusters. Cells were ordered across the trajectory and pseudotime was calculated. DEGs over pseudotime were identified and clustered by their pseudotime expression patterns. For differential expression between branches A and B in Aicda^Cre/−^ mice, BEAM function in Monocle package was used. The top 500 DEGs were used for the branched-heatmap.

### BCR assignment

FASTQ files were preprocessed using the “cellranger vdj” command from 10x Genomics Cell Ranger v3.1.0 for alignment against the Cell Ranger mouse mm10 vdj reference (v3.1.0 July 24, 2019). The resulting FASTA file was split by sample using HTO data and output files were converted to Change-O format. Downstream processing and analyses were performed following the Immcantation pipeline (Gupta et al., 2015). V and J gene usage was determined using IgBlast on the IMGT database and sequences annotated as ‘non-functional’ by IgBLAST were removed from further analysis. Clonal group assignment of sequences was made according to the following requirements: identical V and J gene usage and identical CDR3 length for the IgH and the IgL genes.

### Clonal analysis

Clonal groups were split when differing IgL V and J genes were identified. Germline sequences were reconstructed for each clonal lineage and silent and non-silent mutations were quantified as deviations from the inferred germline. Clone sizes were determined with “countClones” function of Alakazam R package (v1.0.2). Clonal analysis was performed for each mouse separately; mice of the same genotype were grouped together after clonal assignment. Clonal overlap among B cells clusters was quantified and plotted with the UpSetR package (Conway et al., 2017) (v1.4.0). Circos plots were generated using pairwise sharing data obtained from UpSetR, with circlize package (Gu et al., 2014) (v0.4.11). Transition probabilities were manually calculated within each transcriptional cluster for each shared clone present in that cluster. The probability of transition from/to the cluster of interest to the rest of the clusters was calculated based on the presence/absence of the same clone in different clusters. Average transition probabilities were depicted. Interaction networks were generated with Cytoscape (Shannon et al., 2003). Only transitions with probabilities greater than 0.1 are shown. Transition probabilities with differences greater than 0.15 between Aicda^Cre/+^ and Aicda^Cre/−^ mice are colored. Lineage trees were built with IgPhyML, which builds maximum likelihood trees with B cell specific models (HLP19 model). Only clones with more than 2 cells were considered and identical sequences were collapsed. Trees were visualized with Alakazam and ape (v5.4-1) R packages.

Mutational load was calculated using the “ObservedMutations” function of SHazaM R package (v1.0.2) by counting the number of nucleotide mismatches from the germline sequence in the heavy chain variable segment leading up to the CDR3.

### Statistical analysis

Statistical analyses were performed using GraphPad Prism 8. Error bars represent standard deviation of the mean. Normality of the data was assessed with the Anderson-Darling test. For data following a normal distribution, two-tailed unpaired Student’s t-test and ANOVA were used for comparing experimental groups (two or more groups, respectively). Fisher and chi-square tests were used for categorical variables. Krustal-Wallis test was used for comparing not normally distributed data. P-values lower than 0.05 were consider statistically significant.

## SUPPLEMENTAL INFORMATION

**Figure S1. Mouse model and experimental design. A.** Genetic model to study the GC reaction with a fluorescent tracer. Aicda-Cre+/ki; R26tdTom+/ki (Aicda^Cre/+^) and Aicda-Cre-/ki; R26tdTom+/ki (Aicda^Cre/−^) mice are shown. **B.** Experimental approach followed for single cell RNA sequencing.

**Figure S2. Cluster analysis. A.** UMAP plot showing transcriptional clusters obtained before MBC sub-clustering. **B.** UMAP plot showing three transcriptionally distinct memory B cell clusters (Mem0, Mem1, Mem2) conforming the memory B cell pool shown in C. **C.** Dotplot depicting the expression levels of the top 10 upregulated genes in Mem0, Mem1 and Mem2 clusters. **D.** UMAP plot showing enrichment score for GC.DZ and GC.LZ signatures. **E.** Barplot showing enrichment scores for PB/PC differentiation signature in transcriptional clusters identified in Figure 1D. **F.** Heatmap showing expression levels of prePB markers in different Immgen datasets.

**Figure S3. SHM and CSR analysis. A.** p-values of Figure 2A. Statistics were calculated with Krustal-Wallis test. **B.** Quantification of the different isotypes within B cell clusters. **C.** Isotype quantification in B cells according to their different mutational load.

**Figure S4. Comparative clonal analysis of Aicda^Cre/+^ and Aicda^Cre/−^ mice. A.** Bar plot depicting the contribution of the different transcriptional clusters in expanded clones with more than 2 cells in Aicda^Cre/+^ (top) and Aicda^Cre/−^ (bottom) mice. **B.** UpSet plots showing quantification of clonal overlap between clusters identified in Fig 1D in Aicda^Cre/+^ (left) and Aicda^Cre/−^ (right). GC.LZ and GC.DZ populations were grouped and shown as GC for the sake of clarity. **C.** Cytoscape representation of all cluster interactions in Aicda^Cre/+^ and Aicda^Cre/−^ immune response, based on their clonal sharing probabilities. Red and orange connecting lines show the most increased cluster relationships in Aicda^Cre/+^ and Aicda^Cre/−^ mice, respectively.

**Table S1.** Differential scRNA-seq gene expression data for B cell clusters and subclusters.

**Table S2.** Sharing of differentially expressed genes among B cell clusters.

**Table S3.** Clonotype analysis in Aicda^Cre/+^ and Aicda^Cre/−^ mice.

