## Supplemental Figures for "Single cell clonal analysis identifies an AID-dependent pathway of plasma cell differentiation"

**Figure S1. Mouse model and experimental design. A.** Genetic model to study the GC reaction with a fluorescent tracer. Aicda-Cre+/ki; R26tdTom+/ki ($Aicda^{Cre/+}$) and Aicda-Cre-/ki; R26tdTom+/ki ($Aicda^{Cre/-}$) mice are shown. **B.** Experimental approach followed for single cell RNA sequencing.

**Figure S2. Cluster analysis. A.** UMAP plot showing transcriptional clusters obtained before MBC sub-clustering. **B.** UMAP plot showing three transcriptionally distinct memory B cell clusters (Mem0, Mem1, Mem2) conforming the memory B cell pool shown in C. **C.** Dotplot depicting the expression levels of the top 10 upregulated genes in Mem0, Mem1 and Mem2 clusters. **D.** UMAP plot showing enrichment score for GC.DZ and GC.LZ signatures. **E.** Barplot showing enrichment scores for PB/PC differentiation signature in transcriptional clusters identified in Figure 1D. **F.** Heatmap showing expression levels of prePB markers in different Immgen datasets.

**Figure S3. SHM and CSR analysis. A.** p-values of Figure 2A. Statistics were calculated with Krustal-Wallis test. **B.** Quantification of the different isotypes within B cell clusters. **C.** Isotype quantification in B cells according to their different mutational load.

**Figure S4. Comparative clonal analysis of $Aicda^{Cre/+}$ and $Aicda^{Cre/-}$ mice. A.** Bar plot depicting the contribution of the different transcriptional clusters in expanded clones with more than 2 cells in $Aicda^{Cre/+}$ (top) and $Aicda^{Cre/-}$ (bottom) mice. **B.** UpSet plots showing quantification of clonal overlap between clusters identified in Fig 1D in $Aicda^{Cre/+}$ (left) and $Aicda^{Cre/-}$ (right). GC.LZ and GC.DZ populations were grouped and shown as GC for the sake of clarity. **C.** Cytoscape representation of all cluster interactions in $Aicda^{Cre/+}$ and $Aicda^{Cre/-}$ immune response, based on their clonal sharing probabilities. Red and orange connecting lines show the most increased cluster relationships in $Aicda^{Cre/+}$ and $Aicda^{Cre/-}$ mice, respectively.

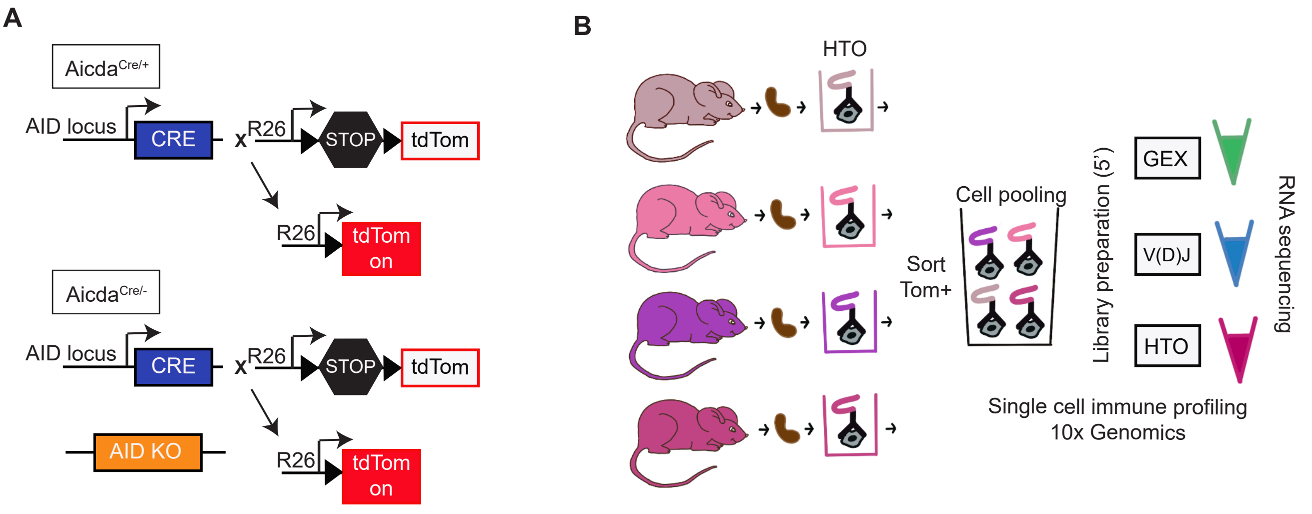
A
Aicda Cre/+
AID locus
CRE
x
R26
STOP
tdTom
R26
tdTom on
Aicda Cre/-
AID locus
CRE
x
R26
STOP
tdTom
AID KO
R26
tdTom on
B
HTO
Cell pooling
Sort Tom+
Library preparation (5')
GEX
V(D)J
HTO
RNA sequencing
Single cell immune profiling
10x Genomics

### Figure S1

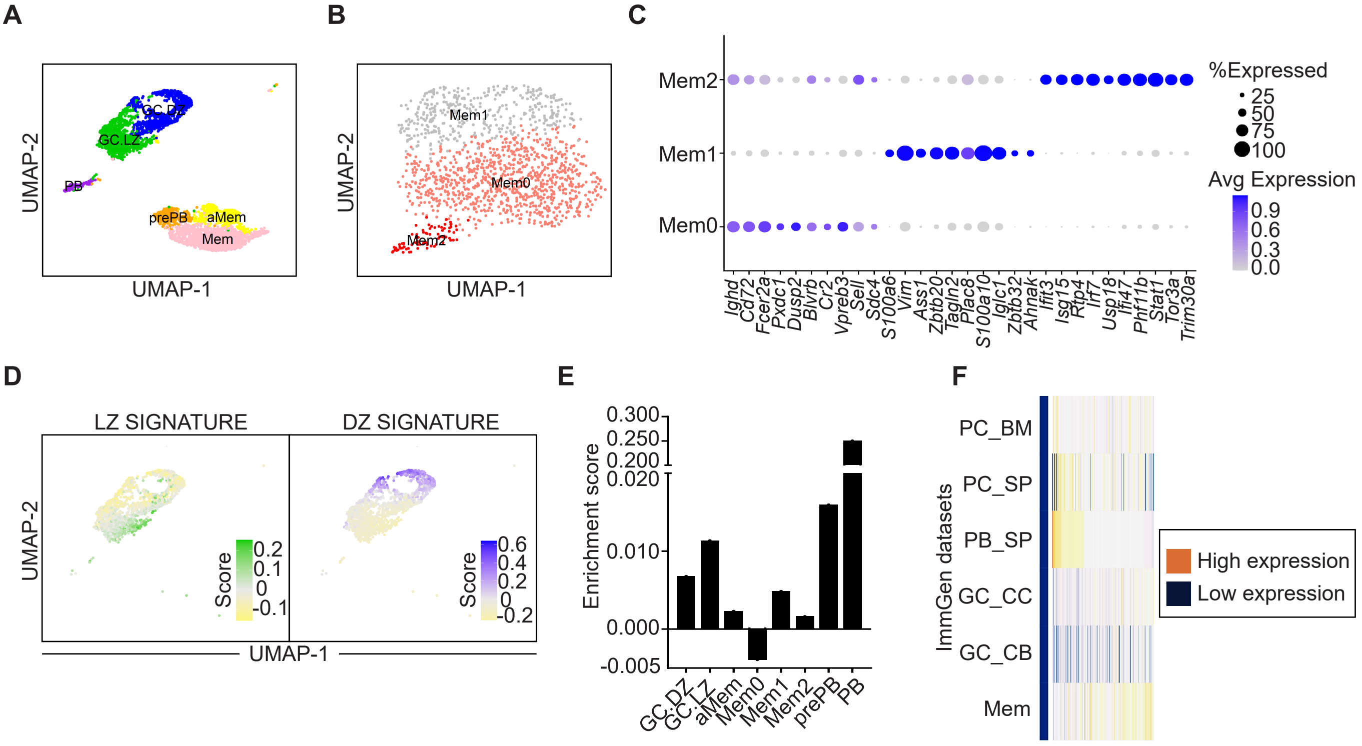
A
UMAP-2
UMAP-1
GC.DZ
GC.LZ
PB
prePB
aMem
Mem
B
UMAP-2
UMAP-1
Mem1
Mem0
Mem2
C
Mem2
Mem1
Mem0
%Expressed
25
50
75
100
Avg Expression
0.9
0.6
0.3
0.0
Ighd
Cd72
Fcer2a
Pxdc1
Dusp2
Blvrb
Cr2
Vpreb3
Sell
Sdc4
S100a6
Vim
Ass1
Zbtb20
Tagln2
Plac8
S100a10
Iglc1
Zbtb32
Ahnak
Ifit3
Isg15
Rtp4
Irf7
Usp18
Ifi47
Phf11b
Stat1
Tor3a
Trim30a
D
LZ SIGNATURE
DZ SIGNATURE
UMAP-2
UMAP-1
Score
0.2
0.1
0
-0.1
0.6
0.4
0.2
0
-0.2
E
Enrichment score
0.300
0.250
0.200
0.020
0.010
0.000
-0.005
GC.DZ
GC.LZ
aMem
Mem0
Mem1
Mem2
prePB
PB
F
ImmGen datasets
PC_BM
PC_SP
PB_SP
GC_CC
GC_CB
Mem
High expression
Low expression

### Figure S2

A

| | GC.LZ | aMem | Mem0 | Mem1 | Mem2 | prePB | PB |
|---|---|---|---|---|---|---|---|
| **GC.DZ** | 0,2321 | **<0,0001** | **<0,0001** | **<0,0001** | **<0,0001** | **<0,0001** | **<0,0001** |
| **GC.LZ** | | **<0,0001** | **<0,0001** | **<0,0001** | **<0,0001** | **<0,0001** | **<0,0001** |
| **aMem** | | | 0,5819 | **<0,0001** | >0,9999 | **0,0083** | 0,1368 |
| **Mem0** | | | | **<0,0001** | >0,9999 | **<0,0001** | **0,0001** |
| **Mem1** | | | | | **0,0003** | >0,9999 | >0,9999 |
| **Mem2** | | | | | | **0,0182** | **0,0394** |
| **prePB** | | | | | | | >0,9999 |

B C

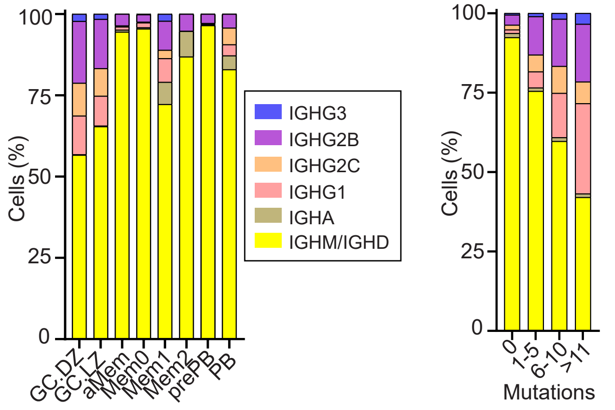


### Figure S3

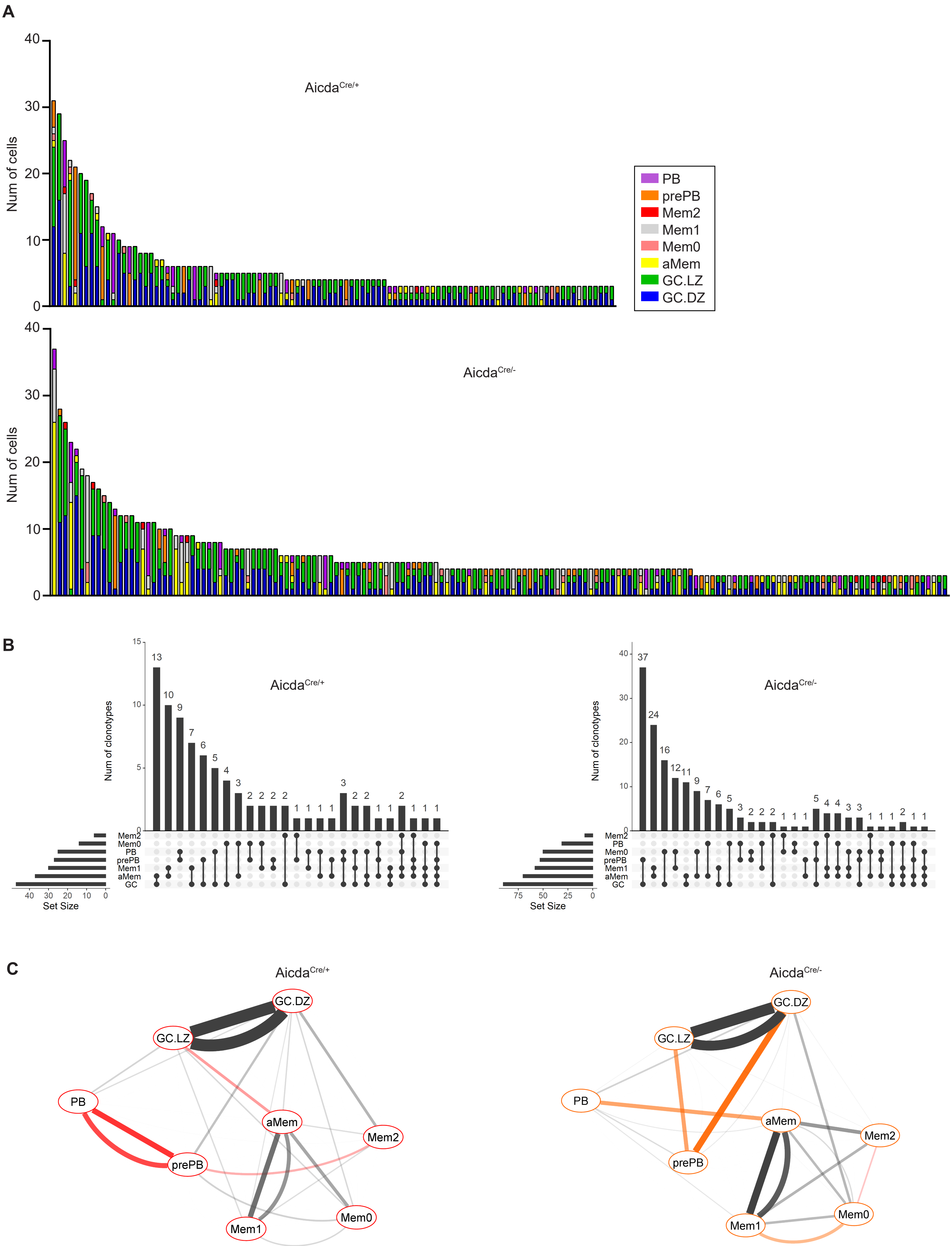
A
Aicda$^{Cre/+}$
Num of cells
PB
prePB
Mem2
Mem1
Mem0
aMem
GC.LZ
GC.DZ
Aicda$^{Cre/-}$
Num of cells
B
Aicda$^{Cre/+}$
Num of clonotypes
Mem2
Mem0
PB
prePB
Mem1
aMem
GC
Set Size
Aicda$^{Cre/-}$
Num of clonotypes
Mem2
PB
Mem0
prePB
Mem1
aMem
GC
Set Size
C
Aicda$^{Cre/+}$
GC.DZ
GC.LZ
PB
aMem
Mem2
prePB
Mem1
Mem0
Aicda$^{Cre/-}$
GC.DZ
GC.LZ
PB
aMem
Mem2
prePB
Mem1
Mem0

**Figure S4**
